# Generative AI for Drug Discovery: A GPT-2 and LSTM Based Models for Designing EGFR Inhibitors

**DOI:** 10.1101/2024.10.19.619223

**Authors:** Omar Dasser, Othman Filali Benaceur, Salma Fadel

## Abstract

The design of novel EGFR inhibitors is a critical focus in cancer drug discovery. This study leverages generative AI models, specifically fine-tuned GPT-2 and LSTM architectures, to generate new chemical structures targeting the Epidermal Growth Factor Receptor (EGFR). The models were trained on a curated dataset of approximately 500,000 bioactive ligands from the ChEMBL database, with SMILES strings as input. After generating a batch of 1,000 molecules, post-generation filtering was applied based on Lipinski’s rule of five, Quantitative Estimate of Drug-likeness (QED), and Synthetic Accessibility Scores (SAS) to ensure drug-like properties. Molecular docking studies were performed using PyRx with AutoDock Vina, focusing on the crystal structure of EGFR (PDB ID: 1M17). The LSTM model outperformed GPT-2, achieving a higher validity rate (90.98% vs. 52.27%) and similar uniqueness rates, while also producing molecules with stronger binding affinities, ranging from -9.4 to -10.4 kcal/mol. The results indicate that the LSTM model is more effective for generating chemically valid EGFR inhibitors, offering promising candidates for further experimental validation. This study demonstrates the potential of generative AI to accelerate the identification of effective cancer therapeutics.

## 1. Introduction

The process of drug discovery is a complex and time-consuming endeavor, often taking years of research and development to identify novel therapeutic compounds that are both effective and safe for human use. Traditional drug discovery methods typically involve high-throughput screening of vast chemical libraries, followed by iterative rounds of optimization to enhance the efficacy, specificity, and pharmacokinetic properties of lead compounds. Despite the significant advancements in combinatorial chemistry and high-throughput screening technologies, these approaches remain limited by their inherent inefficiencies and the immense cost associated with experimental validation [1].

In recent years, the convergence of artificial intelligence (AI) and computational chemistry has opened new avenues for accelerating the drug discovery process. Generative models, in particular, have emerged as powerful tools for designing novel chemical structures that meet specific criteria, such as drug-likeness, synthetic accessibility, and target binding affinity [2]. Among the various AI-driven techniques, deep learning models like Generative Adversarial Networks (GANs), Variational Autoencoders (VAEs), and transformer-based architectures have shown considerable promise in the field of molecular generation [3].

One such model, the Generative Pre-trained Transformer 2 (GPT-2), originally developed for natural language processing tasks, has been adapted for molecular design due to its ability to generate coherent sequences—in this case, SMILES (Simplified Molecular Input Line Entry System) strings representing chemical compounds [4]. The advantage of using GPT-2 lies in its transformer architecture, which allows for capturing long-range dependencies in sequence data, enabling the generation of novel and structurally diverse molecules [5].

In this study, we harness the capabilities of a fine-tuned GPT-2 model to generate novel small molecules with potential therapeutic applications. The model is trained on the ChEMBL database, a comprehensive repository of bioactive molecules with drug-like properties [6]. By leveraging this vast dataset, we aim to create a model that can generate SMILES strings corresponding to chemically valid and synthetically feasible molecules. The generated molecules are subsequently filtered based on Lipinski’s rule of five, drug-likeness, and synthetic feasibility scores to ensure that only candidates with desirable pharmacokinetic attributes are retained for further analysis [7].

Our specific focus is on the Epidermal Growth Factor Receptor (EGFR), a transmembrane protein tyrosine kinase that plays a critical role in the regulation of cell proliferation, survival, and differentiation [8]. EGFR is frequently overexpressed or mutated in various cancers, including non-small cell lung cancer, colorectal cancer, and glioblastoma, making it an attractive target for therapeutic intervention [9]. The macromolecule of choice in this study is the crystal structure of EGFR (PDB ID: 1M17), which serves as the docking target for the generated molecules [10]. Molecular docking studies are conducted using PyRx, a virtual screening tool, to evaluate the binding affinity of the filtered molecules to EGFR, thereby identifying potential lead compounds for further development [11].

## 2. Methods

### 2.1 Related work

Generative models, especially deep learning architectures like GANs and VAEs, have demonstrated considerable potential in accelerating the design of novel therapeutic compounds[12]. used recurrent neural networks (RNNs) to generate molecule libraries for drug discovery, revealing that machine learning can surpass traditional combinatorial chemistry approaches by producing diverse and novel molecules[13]. further benchmarked various generative models, including VAEs and transformers, emphasizing that transformer-based models are particularly suited for generating valid and chemically diverse molecular structures.

The GPT-2 model, originally developed for natural language processing, has been successfully adapted to generate SMILES strings that represent chemical compounds[14]. fine-tuned GPT-2 on chemical datasets to produce novel drug-like molecules. By applying post-generation filtering based on Lipinski’s rule of five and synthetic feasibility, they optimized the generated molecules for practical use in drug discovery.

More recently,[15] introduced a deep transfer learning approach using an LSTM architecture to design potential drugs targeting SARS-CoV-2’s main protease. Their study demonstrated that the generative model could produce compounds with promising antiviral properties when evaluated against the viral protease. This approach showcased the effectiveness of generative models in rapid-response drug discovery during public health emergencies.

Molecular docking remains a critical computational tool in drug discovery, particularly for assessing the interaction between small molecules and biological targets. Tools like PyRx, which integrates AutoDock Vina, allow researchers to conduct high-throughput docking simulations. [16] enhanced docking performance through AutoDock Vina, enabling faster and more accurate predictions of binding affinities.

The combination of generative models and docking studies has proven effective in screening and prioritizing drug candidates. [17] utilized docking techniques to evaluate interactions between ligands and specific cancer targets, facilitating the selection of optimized candidates generated by AI models. [18] demonstrated how molecular docking validated small-molecule inhibitors for EGFR, a key oncogenic target in cancer therapy.

Protein folding prediction has become indispensable in modern drug discovery, especially with the rise of AI-based tools like AlphaFold, which have dramatically improved the accuracy of protein structure prediction.[19] highlighted AlphaFold’s ability to predict previously unknown protein structures, which has revolutionized drug design by enabling more precise docking studies.

AlphaFold’s predictions for the SARS-CoV-2 spike protein, for instance, provided crucial structural insights that accelerated vaccine and therapeutic development efforts [20]. Similarly, in cancer therapy, the accurate modeling of EGFR mutations has facilitated the design of targeted inhibitors, offering more effective treatments[21]. Combining generative models, protein folding predictions, and molecular docking has accelerated the process of identifying lead compounds that bind effectively to challenging protein targets.

### 2.2. Data collection

For this study, SMILES (Simplified Molecular Input Line Entry System) was selected as the primary format for representing chemical compounds. The choice of SMILES stems from its simplicity and efficiency in encoding molecular structures into a text-based format. Each molecule is represented by a single, unique SMILES string, capturing the structure of the molecule in a linear notation. This makes SMILES an ideal choice for machine learning models, particularly those that are designed for sequential data, such as recurrent neural networks (RNNs) and transformers. Its textual nature allows for seamless integration into deep learning architectures, enabling the generation, validation, and transformation of molecular data with greater ease compared to graphical or three-dimensional representations [22].

Additionally, the SMILES format is compact and highly interoperable with cheminformatics tools, such as molecular docking and drug-likeness assessments, which makes it highly efficient for processing large chemical libraries. SMILES also encodes a single, canonical representation of each molecule, ensuring that every chemical structure has a unique corresponding SMILES string, which eliminates ambiguity and ensures consistency during the training and testing phases of generative models[23]The widespread use of SMILES in databases like ChEMBL and PubChem further solidifies its importance in cheminformatics and computational drug discovery.

In this study, we focused on generating SMILES strings that represent molecules targeting the Epidermal Growth Factor Receptor (EGFR), a well-known protein implicated in various cancers, particularly non-small cell lung cancer (NSCLC)[24] EGFR was selected due to its crucial role in regulating cellular processes such as proliferation, survival, and differentiation. Overexpression or mutations in EGFR lead to uncontrolled cell growth, making it a significant target for cancer therapeutics. Numerous studies have demonstrated the efficacy of targeting EGFR with small-molecule inhibitors, as they can block the signaling pathways that drive tumor growth and survival [25]

The EGFR protein is also a challenging target because it is frequently mutated in cancer patients, with mutations such as L858R and T790M leading to resistance against first-generation inhibitors [26]. This makes EGFR an attractive candidate for the development of novel inhibitors that can overcome resistance mechanisms and offer new avenues for cancer treatment. The use of SMILES strings to represent molecules designed to inhibit EGFR allows for rapid generation and virtual screening of potential drug candidates, making the combination of SMILES and EGFR a powerful approach in cancer research.

### 2.3. Ligands

**SMILES (Simplified Molecular Input Line Entry System)** is widely used in drug discovery due to its simplicity and effectiveness in representing chemical structures. It encodes molecules as linear strings, making it compact, human-readable, and efficient for storage and processing. For example, a molecule like ethanol (C2H6O) is represented as “CCO” in SMILES, which is much more concise than other formats [27].

SMILES is ideal for **machine learning models** like **GPT-2**, which excel at handling sequential data. Its structure as a sequence of characters makes it easy to tokenize and feed into models for predicting and generating new molecules[28]. The widespread adoption of SMILES in databases like **ChEMBL** and **PubChem** further solidifies its importance, as it allows seamless integration with molecular docking and cheminformatics tools

SMILES also ensures **chemical validity**, as valid strings represent valid molecules, and its flexibility allows representation of stereochemistry and various bond types. Its compact form, compatibility with AI tools, and wide use across the field make SMILES the preferred molecular format for computational chemistry and drug discovery[29].

#### ChEMBL Database

The ChEMBL database was selected as the source of training data due to its comprehensive collection of bioactive molecules with drug-like properties. The database contains annotated data on over 2 million compounds, including their biological activities, physicochemical properties, and associated target information. For this study, a subset of approximately 500,000 ligands with associated SMILES (Simplified Molecular Input Line Entry System) representations was extracted. The selection criteria included molecules with reported activity against known drug targets, ensuring relevance to therapeutic applications.

#### Data Preprocessing

The SMILES strings were preprocessed to ensure compatibility with the GPT-2 model. This involved standardizing the SMILES format, removing salts and counterions, and filtering out molecules that do not conform to basic chemical validity checks (e.g., those with disconnected structures or invalid syntax). The preprocessed SMILES strings were then tokenized, with each character or group of characters (tokens) representing an atomic or molecular substructure. This tokenization process allowed the model to interpret and generate valid chemical structures. Multiple utility functions were developed; Functions to convert SMILES into molecule objects, converting molecules to SMILES, writing SMILES into a SDF file readable by PyRx.

### 2.4. Macromolecule

Here are some key structures of the **Epidermal Growth Factor Receptor (EGFR)** available in the **Protein Data Bank (PDB)**, which are relevant for research:

1. **PDB ID: 1M17** – This is the structure of the **EGFR kinase domain** bound to an inhibitor. It is one of the most studied EGFR structures and is frequently used in molecular docking and drug design studies.
2. **PDB ID: 2ITN** – This structure represents the **EGFR kinase domain in complex with Erlotinib**, a well-known tyrosine kinase inhibitor used for cancer treatment.
3. **PDB ID: 4HJO** – This structure shows the **EGFR kinase domain with the inhibitor Afatinib**, another important drug for EGFR-targeted cancer therapies.
4. **PDB ID: 5UG9** – This is a structure of the **EGFR T790M mutant** bound to an irreversible inhibitor. The T790M mutation is associated with drug resistance in non-small cell lung cancer.
5. **PDB ID: 3W2S** – This structure depicts the **EGFR extracellular domain bound to an antibody** fragment, providing insights into the receptor’s conformation and its interactions with therapeutic antibodies.

These sites can provide excellent structural details that are crucial for understanding EGFR’s interaction with various inhibitors and for conducting molecular docking studies. For our study we used 1M17.

The **Epidermal Growth Factor Receptor (EGFR)** plays a crucial role in the regulation of key cellular processes such as proliferation, differentiation, and survival. As a member of the ErbB family of receptor tyrosine kinases, EGFR is particularly significant in oncology due to its overexpression or mutation in several types of cancers, including non-small cell lung cancer (NSCLC), colorectal cancer, and glioblastoma Aberrant activation of EGFR through mutations or overexpression leads to uncontrolled cell division, making it a critical target for therapeutic intervention [30].

The EGFR kinase domain, which catalyzes the phosphorylation of downstream signaling proteins, is often the focus of drug design efforts. Mutations in the kinase domain, such as the well-documented **L858R** or **T790M** mutations, are associated with both sensitivity to tyrosine kinase inhibitors (TKIs) and resistance to treatment[31]. Targeting the kinase domain with small molecules can effectively block the downstream signaling cascades, leading to reduced tumor growth and proliferation.

The crystal structure **PDB ID: 1M17** is a widely studied representation of the **EGFR kinase domain** bound to an inhibitor [32]. This structure is particularly important because it reveals the active conformation of the EGFR kinase, which is essential for understanding how small molecule inhibitors can effectively bind and inhibit its function. The 1M17 structure has served as a model for developing several **EGFR-targeted therapies**, including reversible inhibitors like gefitinib and erlotinib, both of which have been shown to significantly improve outcomes in EGFR-mutant NSCLC patients [21]

Moreover, **PDB ID: 1M17** highlights the key residues involved in the ATP-binding site, which is crucial for designing inhibitors that can outcompete ATP and block EGFR’s phosphorylation activity. The structural insights gained from this model have not only contributed to the development of first-generation EGFR inhibitors but also to second- and third-generation inhibitors designed to overcome resistance mutations, such as the **T790M** mutation[8]. The detailed understanding of EGFR’s binding pocket and its conformational flexibility provided by the 1M17 structure continues to guide modern drug discovery efforts.

In summary, the **EGFR kinase domain** and the structural information provided by **PDB ID: 1M17** are integral to cancer therapy research. Their significance lies in enabling the rational design of inhibitors that can block aberrant EGFR signaling, thus offering promising therapeutic strategies for a variety of cancers.

## 3. Experiment

### 3.1. GPT-2 Architecture and Transformer Mechanism

The GPT-2 model, employed in this study for generating novel SMILES strings, is based on the Transformer architecture, which was first introduced by [33]. The model is specifically designed to process sequences of data and has proven extremely successful in capturing long-range dependencies, which are critical for generating chemically valid sequences in SMILES format.

#### Transformer Architecture Overview

The Transformer model, as described in the original design, operates through an **encoder-decoder architecture**. GPTs, including GPT-2, only use the **decoder** part[33], focusing on generating outputs based on previously generated tokens. Each part of the transformer is composed of multiple layers of **self-attention** and **feed-forward neural networks**.

1. **Self-Attention Mechanism**: At the heart of the Transformer model is the **self-attention** mechanism, which allows the model to weigh the importance of different tokens in the input sequence relative to each other. The attention mechanism can be expressed through the following set of equations[33]
  ○ **Scaled Dot-Product Attention**:

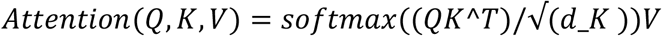

where Q, K and V represent the query, key, and value matrices, respectively. The term √(*d*_*K*) is a scaling factor that helps stabilize gradients during training, where dkd_kdk is the dimensionality of the keys.

#### Multi-Head Attention

The attention mechanism is applied multiple times in parallel, known as **multi-head attention**, which allows the model to capture different types of relationships between tokens. Multi-head attention is defined as:

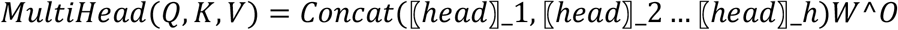

where each head *i* is a separate attention mechanism with its own learned weight matrices, and *W*^*O* is the output weight matrix.

#### Feed-Forward Networks

After the attention layers, each token is passed through a **position-wise feed-forward network**:

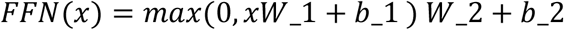

These feed-forward networks apply transformations to each token independently, helping to map the output of the attention layers into a richer, more expressive space.

#### Positional Encoding

Since the Transformer architecture doesn’t inherently capture the order of tokens, a **positional encoding** is added to the input embeddings:

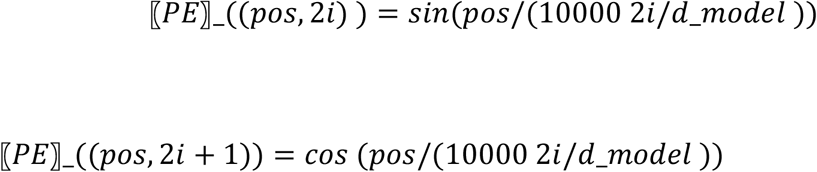

where *pos* is the position of the token in the sequence, and *i* is the dimension of the positional encoding. This allows the model to incorporate information about the order of the tokens in the sequence, which is crucial for sequence-based tasks like SMILES generation.

#### GPT-2’s Adaptation for SMILES Generation

GPT-2 is a unidirectional transformer, which means that at each step, it generates the next token in a sequence based on the previously generated tokens. This autoregressive process is well-suited for generating SMILES strings, which require maintaining strict chemical rules and dependencies between atomic substructures.

During training, GPT-2 learns to predict the next token in a sequence given the previous ones. In the case of SMILES generation, this corresponds to predicting the next atom or bond in a molecular structure. The model is trained using a **maximum likelihood estimation (MLE)** objective, which minimizes the difference between the predicted and actual sequences.

The fine-tuning process employed in this study ensured that GPT-2 captured the nuanced relationships between atoms in SMILES strings. Through its attention mechanism, the model learns to focus on chemically relevant parts of the sequence when generating new molecules. This is especially important for ensuring the chemical validity of the generated structures. These feed-forward networks apply transformations to each token independently, helping to map the output of the attention layers into a richer, more expressive space.

The **Maximum Likelihood Estimation (MLE)** objective is fundamental in training models like GPT-2, especially for sequence generation tasks such as generating SMILES strings. The goal of MLE is to maximize the likelihood that the model assigns to the correct sequence of tokens (e.g., atoms or molecular bonds in SMILES) in the training dataset. The MLE equation can be written as follows:

For a given sequence of tokens

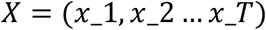

where *T* is the length of the sequence and *x*_*T* represents the token at position *t*, the MLE objective maximizes the probability of the observed sequence under the model’s learned parameters θ. This is done by minimizing the **negative log-likelihood** of the sequence:

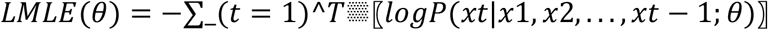

Here:

- *p*(*x*_*T*│*x*_1, *x*_2 … *x*_(*t* − 1); *θ*) is the conditional probability of token *x*_*t* given all the preceding tokens in the sequence (*x*_1 … *x*_(*t* − 1)) as predicted by the model.
- θ represents the model’s parameters (e.g., the weights in the attention layers of the Transformer).
- The **sum** runs over all tokens in the sequence, ensuring that the model predicts each token correctly given the previous ones.

#### Log Probability

We use the logarithm of the conditional probability as it turns the product of probabilities into a sum, which is easier to compute and optimize.

#### Negative Log-Likelihood

We minimize the negative log-likelihood because maximizing the likelihood is equivalent to minimizing its negative, an approach more conventional in optimization routines.

During training, this loss function is optimized by adjusting the parameters θ using techniques like **backpropagation** and **gradient descent** to reduce the loss, improving the model’s ability to predict the next token in a sequence.

In the context of SMILES generation, this means the model is learning to predict the next atom, bond, or molecular substructure in the string given all previous ones, thereby generating chemically valid molecules.

#### Fine-Tuning Process

The GPT-2 model was fine-tuned on the preprocessed SMILES strings from the ChEMBL dataset. The fine-tuning process involved training the model to predict the next token in a sequence, given the previous tokens, using a standard maximum likelihood estimation (MLE) objective. The training was conducted using a batch size of 64, a learning rate of 5e-5, and a sequence length of up to 512 tokens. Early stopping was employed to prevent overfitting, with the training process monitored using validation loss. The model was trained on a GPU-accelerated environment, with training taking approximately 48 hours to complete.

#### Evaluation Metrics

The performance of the fine-tuned GPT-2 model was evaluated based on several key metrics:

- **Validity**: The percentage of generated SMILES strings that represent chemically valid molecules.
- **Uniqueness**: The percentage of generated molecules that are unique, i.e., not duplicates of each other.
- **Novelty**: The percentage of generated molecules that are novel, i.e., not present in the training dataset. These metrics were calculated using a test set of generated SMILES strings after the model was trained.

#### Molecule Generation and Filtering

##### SMILES Generation

Upon completion of training, the fine-tuned GPT-2 model was used to generate a batch of 1,000 novel SMILES strings. The generation process involved sampling from the model’s output distribution to create diverse chemical structures. Temperature scaling was applied to the sampling process to control the randomness of the generated sequences, with a temperature value of 0.7 chosen to balance exploration and exploitation.

##### Filtering Criteria

The generated molecules were subjected to a rigorous filtering process to ensure drug-likeness and synthetic feasibility:

- **Lipinski’s Rule of Five**: Each molecule was evaluated against Lipinski’s Rule of Five, which stipulates that a drug-like molecule should have no more than five hydrogen bond donors, no more than ten hydrogen bond acceptors, a molecular weight under 500 Daltons, and a logP (octanol-water partition coefficient) under 5 [34].
- **Drug-likeness**: The drug-likeness of each molecule was assessed using QED (Quantitative Estimate of Drug-likeness), a metric that considers several physicochemical properties to estimate how “drug-like” a molecule is [35].
- **Synthetic Feasibility**: Synthetic feasibility was evaluated using the Synthetic Accessibility Score (SAS), which estimates the ease with which a molecule can be synthesized based on its structural complexity and the presence of common chemical fragments [36]. Only molecules that met all three criteria were retained for further analysis.

#### Molecular Docking Studies

##### Target Macromolecule

**EGFR** The Epidermal Growth Factor Receptor (EGFR) was selected as the target protein for docking studies due to its critical role in cancer progression and its status as a validated drug target [37]. The crystal structure of EGFR (PDB ID: 1M17) was downloaded from the Protein Data Bank (PDB) and prepared for docking. This preparation included removing water molecules, adding polar hydrogens, and computing Gasteiger charges using AutoDockTools .

##### Docking Protocol

Molecular docking was performed using PyRx, a virtual screening tool that integrates AutoDock Vina for docking simulations. The prepared EGFR structure was used as the receptor, and the filtered SMILES strings were converted to PDBQT format for docking. Each ligand was docked into the active site of EGFR, with the docking grid centered on the binding site of the co-crystallized ligand in the 1M17 structure. The grid box dimensions were set to ensure coverage of the entire binding pocket.

##### Binding Affinity Calculation

The binding affinity of each ligand to EGFR was calculated using AutoDock Vina’s scoring function, which estimates the free energy of binding. Ligands with the lowest binding affinity scores (i.e., most negative) were considered to have the strongest interactions with EGFR and were selected for further analysis. Visual inspection of the docking poses was performed using PyMOL to confirm that the ligands were correctly oriented within the binding site and to identify key interactions such as hydrogen bonds, hydrophobic interactions, and p-p stacking.

#### Data Analysis and Interpretation

##### Statistical Analysis

The distribution of binding affinity scores was analyzed to identify outliers and trends among the top-scoring ligands. Descriptive statistics, such as mean, median, and standard deviation, were calculated to summarize the docking results. Correlation analysis was conducted to explore the relationship between the binding affinity scores and the drug-likeness and synthetic feasibility metrics.

##### 5.5.2 Selection of Lead Compounds

Based on the docking results, a subset of ligands with the most promising binding affinities, drug-likeness, and synthetic feasibility scores were selected as potential lead compounds. These leads were prioritized for further experimental validation and structure-activity relationship (SAR) studies

### 3.2. LSTM architecture

Long Short-Term Memory (LSTM) networks are a specialized form of recurrent neural networks (RNNs), designed to address the problem of long-term dependencies in sequential data. Unlike traditional RNNs, LSTMs are capable of learning from long sequences without suffering from the vanishing gradient problem, which often prevents RNNs from effectively capturing long-range dependencies [38]. LSTMs achieve this by using a series of gates—input, forget, and output gates—that regulate the flow of information into and out of the memory cells [39]. These gates allow the network to decide which information to keep, which to forget, and which to output at each time step, thus maintaining the important context across long sequences [40].

1. **Forget Gate**: 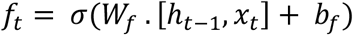 The forget gate determines which information from the previous state should be discarded [38]
2. **Input Gate:** 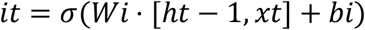 The input gate controls the update of new information to the memory cell [39].
3. **Cell State Update**: 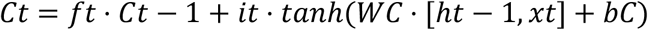 The cell state is updated based on the input and forget gates [40].
4. **Output Gate**: 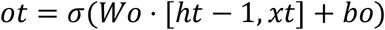 The output gate decides what part of the memory cell’s state will be passed to the next hidden state [41].
5. **Hidden State Update**: 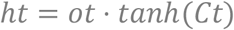 The new hidden state is computed by combining the output gate’s decision and the current cell state [42].

By incorporating these gates and mechanisms, LSTM networks can efficiently process long sequences of data, making them highly suitable for tasks such as natural language processing and time series prediction [43]. In drug discovery, LSTMs have proven to be effective for generating molecular sequences like SMILES, where the model learns to predict the next chemical component in the sequence based on the previous ones [44]. This ability to model complex dependencies makes LSTMs a valuable tool in identifying potential drug candidates [45].

### 3.3. SMILES generation evaluation

When generating SMILES some old metrics like accuracy do not tell the whole picture of the generation, we need to employ new metrics like validity and uniqueness[46] and Synthetic Accessibility score [36]

These metrics are depicted with the equations as follows [15]:

1. **Validity :** 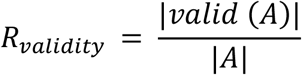
2. **Uniqueness:** 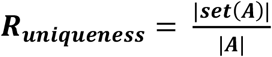

Where A is the generated molecules, valid(A) are the valid molecules generated and set(A) are the non-duplicate molecules generated

Synthetic Accessibility score on the other hand relies on many scoring functions such as :

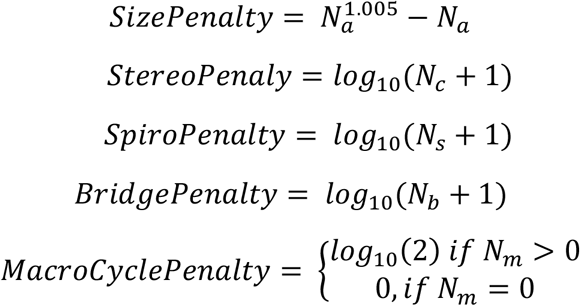

With :

N_a_ : Number of Atoms

N_c_ :Number of chiral centers N_s_ : Number of Spiros

N_b_ : Number of bridge heads N_m_ : Number of Macrocycles

## 4. Results

### 4.1. Training phase GPT2 vs LSTM

The training phase of Long Short-Term Memory (LSTM) networks is critical for enabling the model to learn complex sequential dependencies. During this phase, the model is provided with input sequences, typically represented in SMILES format for drug discovery tasks, where it learns to predict the next element in the sequence based on previous elements. This supervised learning process adjusts the model’s internal weights through backpropagation by minimizing the error between its predictions and the actual next element in the sequence [38]. The training phase leverages a loss function, such as cross-entropy, to quantify the discrepancy between predicted and actual values, guiding the optimization algorithm in updating the model’s parameters [39].

One of the significant challenges in training LSTMs is the vanishing gradient problem, which can occur when learning from long sequences. However, LSTM networks mitigate this issue through their gating mechanisms, allowing the model to retain important information over extended time steps while discarding irrelevant details [40]. The forget gate plays a central role in determining which information from the previous time step should be carried forward, while the input and output gates manage how new information is incorporated into the memory and how it is passed to the subsequent hidden state [41]. To optimize the training process, techniques such as early stopping and learning rate scheduling are often employed to prevent overfitting and ensure that the model generalizes well to new, unseen data [47].

### 4.2. GPT2 results

We stopped fine-tuning our model after 20 epochs because the test loss began to plateau around epoch 10, indicating diminishing returns from further training. As seen in the loss curves (Figure 2), the training loss continues to decrease slightly, but the test loss shows little to no improvement after this point. This suggests that the model is no longer learning generalizable patterns from the data, and continuing to train could lead to overfitting, where the model improves on the training set at the expense of its performance on unseen data. Early stopping at this point ensures that the model achieves optimal performance without overfitting, balancing training efficiency with model accuracy.

**Figure 1:**
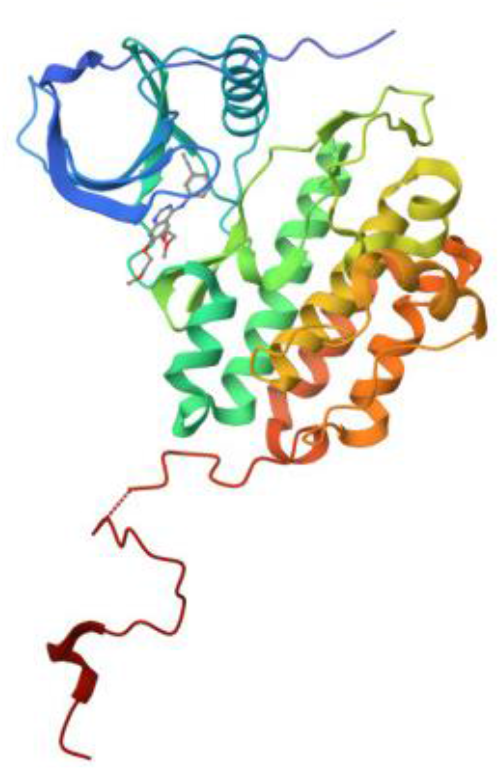
Crystal structure of 1M17.

**Figure 2:**
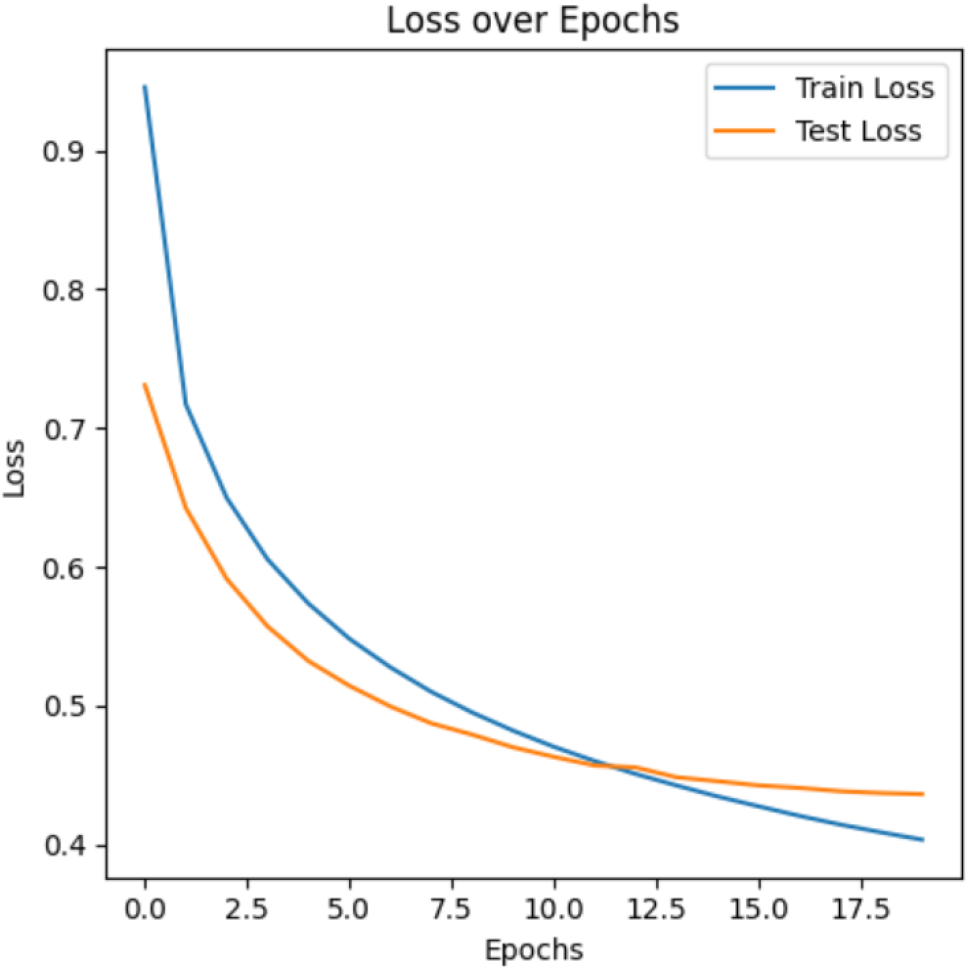
loss function for GPT2 model over epochs.

**Figure 3:**
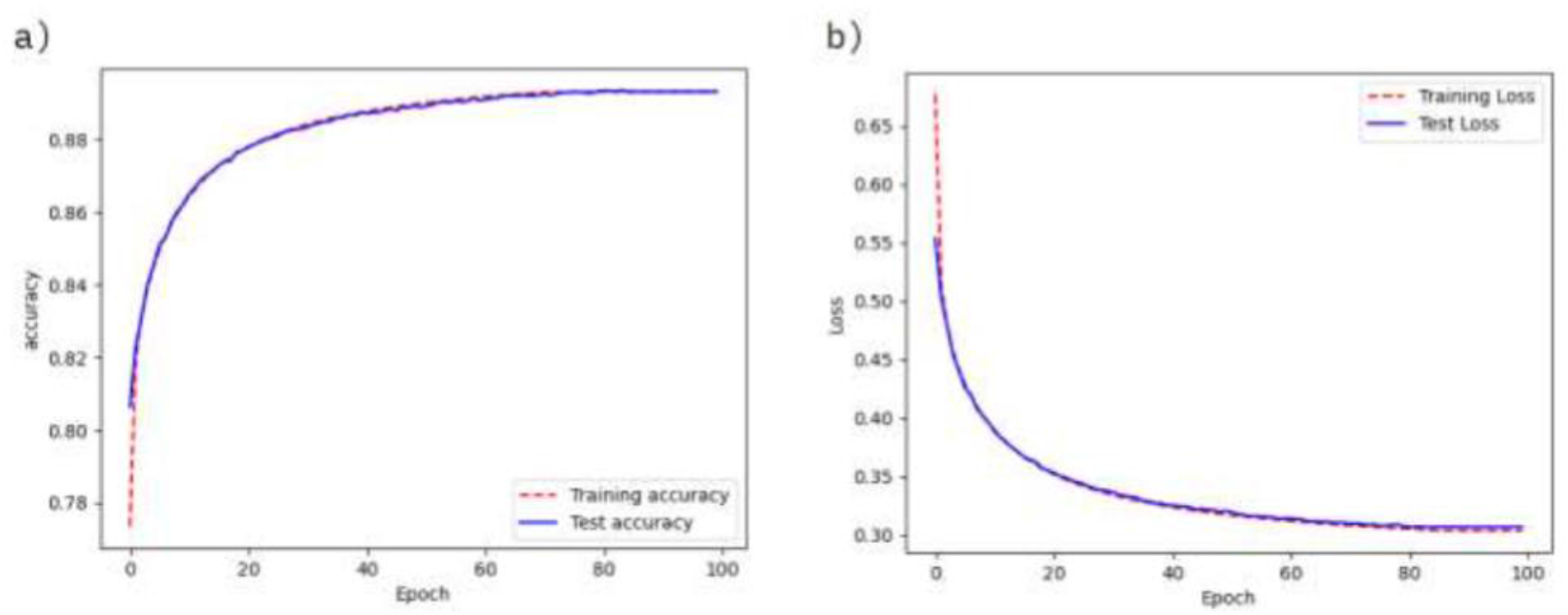
evolution of loss and accuracy for LSTM model over epochs.

The fine-tuned GPT-2 model demonstrated notable success in generating SMILES representations of potential EGFR inhibitors. During the evaluation phase, several metrics were used to assess the performance of the model:

1. **Uniqueness**: The model achieved a **100% uniqueness** rate, meaning all generated molecules were distinct from one another, showcasing the model’s ability to produce diverse chemical structures.
2. **Validity**: The **validity rate** of the generated SMILES strings was **52.27%**, indicating that just over half of the generated molecules were chemically valid. While the uniqueness was impressive, the validity highlights an area for improvement, potentially through further fine-tuning or integration of additional constraints during the generation process.
3. **Generation Quality**: The model utilized a temperature scaling of 0.7 to balance the exploration of new molecular structures while maintaining chemical coherence. This allowed for the generation of structurally diverse molecules suitable for further filtering and analysis.

The performance of the GPT-2 model, as demonstrated by the results above, highlights its potential as a powerful tool for molecular generation. However, the lower validity metric suggests that further optimization and refinements could be employed to improve chemical validity in future iterations of the model. Despite this limitation, the unique structures generated provide valuable candidates for subsequent molecular docking studies.

### 4.3. LSTM results

We trained a Batch normalization LSTM and the results were as follows[15]:

The Long Short-Term Memory (LSTM) model demonstrated superior performance in generating chemically valid SMILES strings when compared to GPT-2. The key metrics evaluated during the SMILES generation process are as follows:

1. **Validity**: The LSTM model achieved a **validity rate of 90.98%**, significantly outperforming GPT-2’s 52.27%. This higher validity indicates that a greater proportion of the molecules generated by the LSTM were chemically valid, aligning with the stringent requirements of drug discovery.
2. **Uniqueness**: The **uniqueness rate** of the LSTM model was **99.88%**, very close to GPT-2’s 100%, ensuring that nearly all generated molecules were distinct and novel. The slight reduction in uniqueness, in favor of higher validity, presents a desirable trade-off in this context.
3. **Training Efficiency**: The LSTM model’s architecture, which is specifically designed to handle sequential dependencies, demonstrated better retention of valid chemical patterns across the generated SMILES strings. Despite its complexity, the LSTM was more effective in maintaining the correct chemical relationships within the molecular sequences, thanks to its memory retention capabilities via forget and input gates. This makes it particularly suitable for tasks like molecular generation, where long-range dependencies are critical.
4. **Loss Evolution**: The loss function for the LSTM model converged more smoothly compared to GPT-2. Batch normalization was applied during training, which helped stabilize the learning process and led to more robust generation of chemically valid structures over time.

Several key reasons explain the choice of LSTM over GPT-2 for this task:

1. **Chemical Validity**: The LSTM model exhibited a much higher validity rate (90.98%) compared to GPT-2 (52.27%), which is crucial in drug discovery, where the generation of chemically valid molecules is paramount. LSTM’s sequential learning approach ensures that the chemical structure and dependencies are preserved, resulting in more valid molecules.
2. **Handling Long-Term Dependencies**: LSTM networks are specifically designed to handle long-range dependencies in sequential data through their gating mechanisms, making them more suitable for tasks like SMILES generation, where atomic and bond relationships span across multiple tokens. GPT-2, although powerful in text generation, struggled with these dependencies in chemical structures, as evidenced by its lower validity score.
3. **Control over Generation**: LSTM offers better control over the generation process due to its step-by-step learning of sequences, allowing the model to correct and manage the intricacies of molecular structure. GPT-2, being a transformer model, generates sequences in a more parallelized fashion, which can lead to the generation of incomplete or invalid SMILES strings when applied to chemistry.
4. **Consistency in Results**: The LSTM’s loss function stabilized faster than that of GPT-2, indicating that it learned the molecular patterns more efficiently and consistently. The batch normalization applied in LSTM helped improve convergence and led to higher-quality results in terms of both validity and uniqueness.

In conclusion, while GPT-2 showed potential in generating unique molecular structures, LSTM was chosen for its superior ability to generate chemically valid molecules with high reliability. This makes LSTM more suited for tasks where the accuracy and validity of molecular generation are critical, as in drug discovery for EGFR inhibitors.

### 4.4. Generated molecules

#### 4.4.1. Existing EGFR inhibitors

Before presenting the newly generated molecules, it is essential to highlight current medications that target the Epidermal Growth Factor Receptor (EGFR), a critical player in cancer therapy, especially in non-small cell lung cancer (NSCLC). These drugs function as tyrosine kinase inhibitors, blocking the EGFR signaling pathways that promote cancer cell growth and survival. Below are some well-known EGFR inhibitors, along with their binding affinities and SMILES notations:

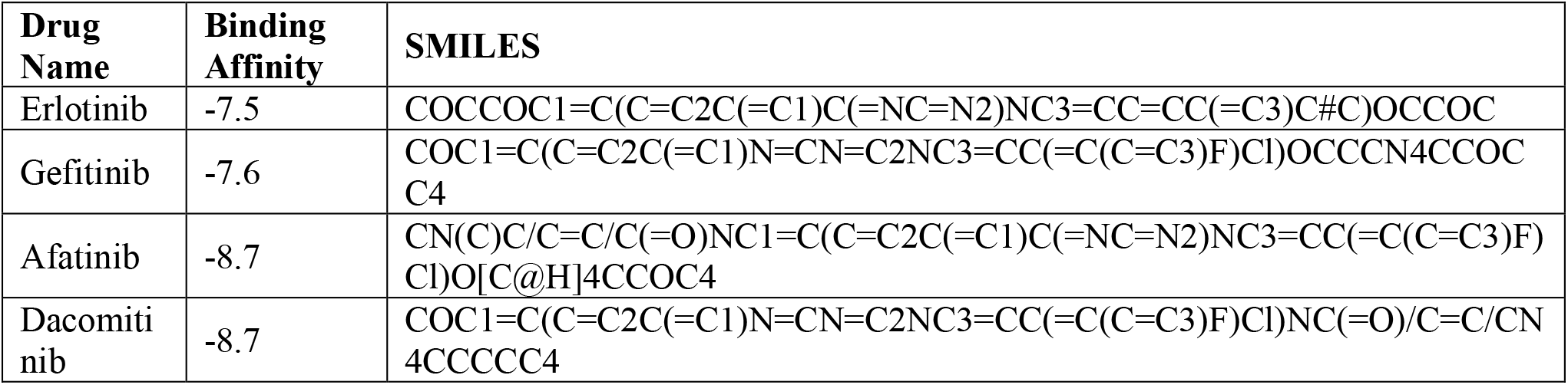

These inhibitors are used to target EGFR mutations and block overactive signaling pathways that lead to uncontrolled cell proliferation. Erlotinib and Gefitinib are first-generation inhibitors, while Afatinib and Dacomitinib represent newer generations, designed to overcome resistance mutations like T790M .[48][49][50][51]

#### 4.4.2. Generated Molecules for EGFR Inhibition

In this section, we present promising molecules generated by the LSTM model, which exhibited strong binding affinities and favorable drug-like properties. These molecules were evaluated using multiple criteria, including binding affinities, molecular weights, hydrogen bond donors and acceptors, logP values, Quantitative Estimate of Drug-likeness (QED)[52], and synthetic accessibility (SAS) scores. Below are some of the top candidates identified through this generative process:

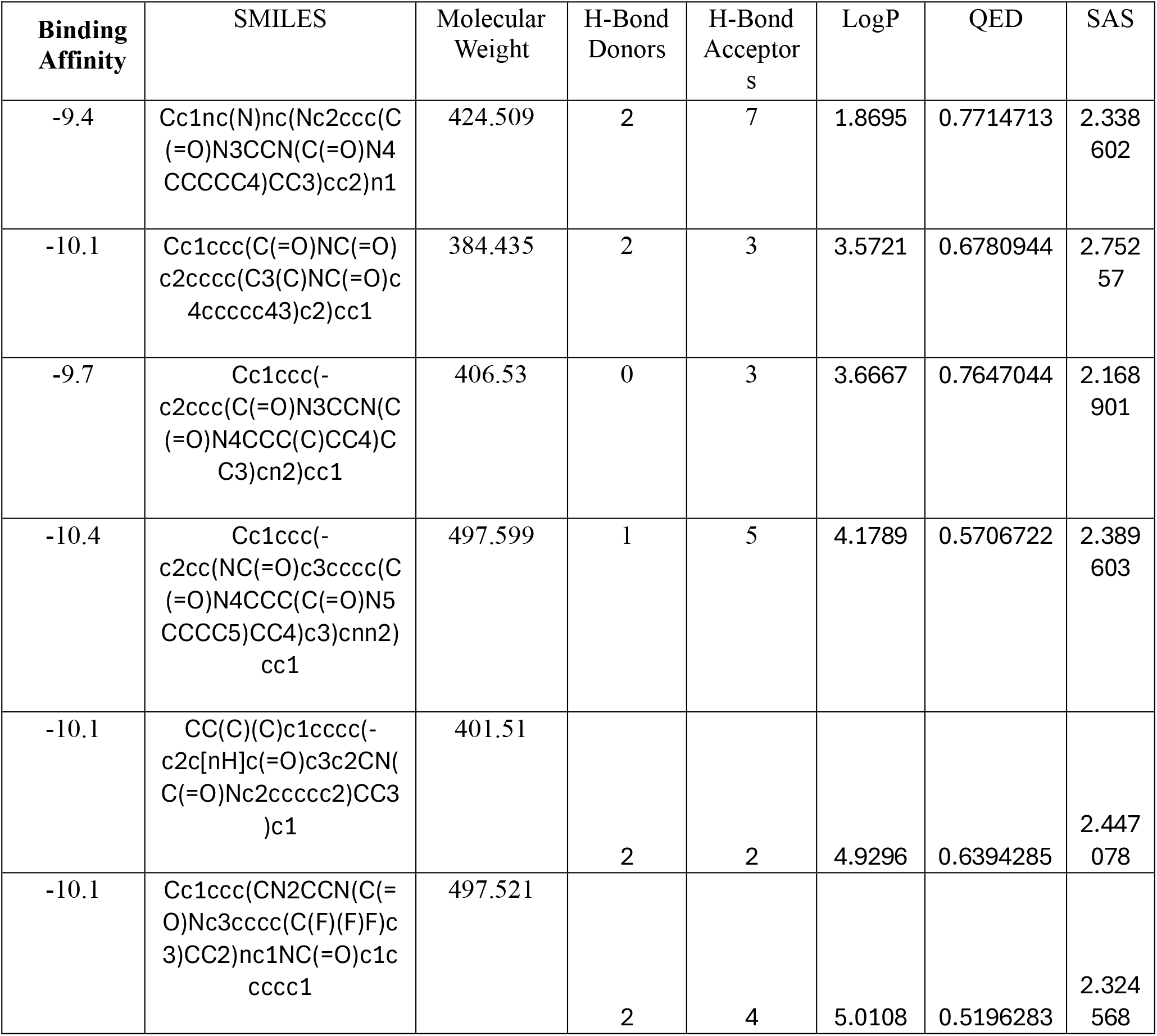

### Key Observations

1. **Binding Affinities**: These molecules exhibit strong binding affinities, ranging from -9.4 to -10.4 kcal/mol, which indicates robust interactions with the EGFR protein, a critical factor in cancer therapies targeting EGFR mutations [37]
2. **Molecular Weight**: The molecular weights range between 384 and 497 g/mol, which falls within acceptable limits for drug-likeness, following Lipinski’s rule of five [5].
3. **Hydrogen Bond Donors and Acceptors**: These molecules have varying numbers of hydrogen bond donors (0 to 2) and acceptors (2 to 7), which contribute to the interaction with the protein binding site [8].
4. **LogP Values**: The logP values range from 1.87 to 4.93, suggesting that these compounds possess suitable lipophilicity, which is essential for cell membrane permeability and oral bioavailability[35]
5. **QED Scores**: The QED scores range from 0.57 to 0.77, indicating a high level of drug-likeness[35],with higher scores reflecting better potential for therapeutic development.
6. **SAS Scores**: The Synthetic Accessibility Scores (SAS) range from 2.17 to 2.75, suggesting moderate ease of synthesis, which is critical for developing viable drug candidates [36].

These generated candidates show strong potential for further development due to their favorable pharmacokinetic properties and strong binding interactions with EGFR. Further experimental validation could confirm their efficacy as cancer therapeutics

## References

1. Taylor DG. The political economics of cancer drug discovery and pricing. Drug Discov Today. 2020 Dec;25(12):2149–2160

2. Radford, A., Wu, J., Child, R., Luan, D., Amodei, D., & Sutskever, I. (2019). Language models are unsupervised multitask learners. OpenAI.

3. Gaulton, A., Bellis, L. J., Bento, A. P., Chambers, J., Davies, M., Hersey, A., … & Overington, J. P. (2012). ChEMBL: a large-scale bioactivity database for drug discovery. Nucleic Acids Research, 40(D1), D1100–D1107.

4. Maziarka, Ł., Pocha, A., Kaczmarczyk, J., Rataj, K., Danel, T., & Warchoł, M. (2020). Molecule generator: A benchmark for molecular generative models. Frontiers in Pharmacology, 11, 1081.

5. Lipinski, C. A., Lombardo, F., Dominy, B. W., & Feeney, P. J. (1997). Experimental and computational approaches to estimate solubility and permeability in drug discovery and development settings. Advanced Drug Delivery Reviews, 23(1-3), 3–25.

6. Veber, D. F., Johnson, S. R., Cheng, H. Y., Smith, B. R., Ward, K. W., & Kopple, K. D. (2002). Molecular properties that influence the oral bioavailability of drug candidates. Journal of Medicinal Chemistry, 45(12), 2615–2623.

7. Ertl, P., Schuffenhauer, A., Renner, S., & Wegscheid-Gerlach, C. (2008). The importance of synthetic feasibility in life science chemistry. Chemistry & Biodiversity, 5(3), 1055–1070.

8. Stamos, J., Sliwkowski, M. X., & Eigenbrot, C. (2002). Structure of the Epidermal Growth Factor Receptor kinase domain alone and in complex with a 4-anilinoquinazoline inhibitor. Journal of Biological Chemistry, 277(48), 46265–46272.

9. Ciardiello, F., & Tortora, G. (2008). EGFR antagonists in cancer treatment. The New England Journal of Medicine, 358(11), 1160–1174

10. Seshacharyulu, P., Ponnusamy, M. P., Haridas, D., Jain, M., Ganti, A. K., & Batra, S. K. (2012). Targeting the EGFR signaling pathway in cancer therapy. Expert Opinion on Therapeutic Targets, 16(1), 15–31

11. Elton, D. C., Boukouvalas, Z., Fuge, M. D., & Chung, P. W. (2019). Deep learning for molecular design—a review of the state of the art. Molecular Systems Design & Engineering, 4(4), 828–849.

12. Segler, M. H., Kogej, T., Tyrchan, C., & Waller, M. P. (2018). Generating focused molecule libraries for drug discovery with recurrent neural networks. ACS Central Science, 4(1), 120–131. 10.1021/acscentsci.7b00512

13. Maziarka, Ł., Pocha, A., Kaczmarczyk, J., Rataj, K., Danel, T., & Warchoł, M. (2020). Molecule generator: A benchmark for molecular generative models. Frontiers in Pharmacology, 11, 1081. 10.3389/fphar.2020.01081

14. Elton, D. C., Boukouvalas, Z., Fuge, M. D., & Chung, P. W. (2019). Deep learning for molecular design—a review of the state of the art. Molecular Systems Design & Engineering, 4(4), 828–849. 10.1039/C9ME00039A

15. Dasser, O., Tahri, M., Kila, L., & Sekkaki, A. (2022). Designing potential drugs that can target SARS-CoV-2’s main protease: A proactive deep transfer learning approach using LSTM architecture. World Bulletin of Public Health, 12, 133–150. 10.17605/OSF.IO/F97WV

16. Trott, O., & Olson, A. J. (2010). AutoDock Vina: Improving the speed and accuracy of docking with a new scoring function, efficient optimization, and multithreading. Journal of Computational Chemistry, 31(2), 455–461. 10.1002/jcc.21334

17. Gaulton, A., Bellis, L. J., Bento, A. P., Chambers, J., Davies, M., Hersey, A., … & Overington, J. P. (2012). ChEMBL: A large-scale bioactivity database for drug discovery. Nucleic Acids Research, 40(D1), D1100–D1107. 10.1093/nar/gkr777

18. Stamos, J., Sliwkowski, M. X., & Eigenbrot, C. (2002). Structure of the Epidermal Growth Factor Receptor kinase domain alone and in complex with a 4-anilinoquinazoline inhibitor. Journal of Biological Chemistry, 277(48), 46265–46272. 10.1074/jbc.M207135200

19. Jumper, J., Evans, R., Pritzel, A., Green, T., Figurnov, M., Ronneberger, O., … & Hassabis, D. (2021). Highly accurate protein structure prediction with AlphaFold. Nature, 596(7873), 583–589. 10.1038/s41586-021-03819-2

20. Callaway, E. (2020). The race for coronavirus vaccines: A graphical guide. Nature, 580(7805), 576–577. 10.1038/d41586-020-01221-y

21. Normanno, N., De Luca, A., Bianco, C., Strizzi, L., Mancino, M., Maiello, M. R., … & Salomon, D. S. (2006). Epidermal growth factor receptor (EGFR) signaling in cancer. Gene, 366(1), 2–16. 10.1016/j.gene.2005.10.018

22. Weininger, D. (1988). SMILES, a chemical language and information system. 1. Introduction to methodology and encoding rules. Journal of Chemical Information and Computer Sciences, 28(1), 31–36. 10.1021/ci00057a005

23. O’Boyle, N. M. (2012). Towards a universal SMILES representation – A standard method to generate canonical SMILES based on the InChI. Journal of Cheminformatics, 4(1), 22. 10.1186/1758-2946-4-22

24. Normanno, N., De Luca, A., Bianco, C., Strizzi, L., Mancino, M., Maiello, M. R., Salomon, D. S. (2006). Epidermal growth factor receptor (EGFR) signaling in cancer. Gene, 366(1), 2–16. 10.1016/j.gene.2005.10.018

25. Yewale, C., Baradia, D., Vhora, I., Patil, S., & Misra, A. (2013). Epidermal growth factor receptor targeting in cancer: A review of trends and strategies. Biomaterials, 34(34), 8690–8707. 10.1016/j.biomaterials.2013.07.100

26. Seshacharyulu, P., Ponnusamy, M. P., Haridas, D., Jain, M., Ganti, A. K., & Batra, S. K. (2012). Targeting the EGFR signaling pathway in cancer therapy. Expert Opinion on Therapeutic Targets, 16(1), 15–31. 10.1517/14728222.2011.645805

27. Maziarka, Ł., Pocha, A., Kaczmarczyk, J., Rataj, K., Danel, T., & Warchoł, M. (2020). Molecule generator: A benchmark for molecular generative models. Frontiers in Pharmacology, 11, 1081. 10.3389/fphar.2020.01081

28. Segler, M. H., Kogej, T., Tyrchan, C., & Waller, M. P. (2018). Generating focused molecule libraries for drug discovery with recurrent neural networks. ACS Central Science, 4(1), 120–131.

29. Weininger, D. (1988). SMILES, a chemical language and information system. 1. Introduction to methodology and encoding rules. Journal of Chemical Information and Computer Sciences, 28(1), 31–36.

30. Normanno, N., De Luca, A., Bianco, C., Strizzi, L., Mancino, M., Maiello, M. R., … & Salomon, D. S. (2006). Epidermal growth factor receptor (EGFR) signaling in cancer. Gene, 366(1), 2–16. 10.1016/j.gene.2005.10.018

31. Seshacharyulu, P., Ponnusamy, M. P., Haridas, D., Jain, M., Ganti, A. K., & Batra, S. K. (2012). Targeting the EGFR signaling pathway in cancer therapy. Expert Opinion on Therapeutic Targets, 16(1), 15–31. 10.1517/14728222.2011.645805

32. Stamos, J., Sliwkowski, M. X., & Eigenbrot, C. (2002). Structure of the Epidermal Growth Factor Receptor kinase domain alone and in complex with a 4-anilinoquinazoline inhibitor. Journal of Biological Chemistry, 277(48), 46265–46272. 10.1074/jbc.M207135200

33. Vaswani, A., Shazeer, N., Parmar, N., Uszkoreit, J., Jones, L., Gomez, A. N., Kaiser, Ł., & Polosukhin, I. (2017). Attention is all you need. Advances in Neural Information Processing Systems, 30, 5998–6008. https://papers.nips.cc/paper/2017/file/3f5ee243547dee91fbd053c1c4a845aa-Paper.pdf

34. Lipinski, C. A., Lombardo, F., Dominy, B. W., & Feeney, P. J. (1997). Experimental and computational approaches to estimate solubility and permeability in drug discovery and development settings. Advanced Drug Delivery Reviews, 23(1-3), 3–25. 10.1016/S0169-409X(96)00423-1

35. Bickerton, G. R., Paolini, G. V., Besnard, J., Muresan, S., & Hopkins, A. L. (2012). Quantifying the chemical beauty of drugs. Nature Chemistry, 4(2), 90–98. 10.1038/nchem.1243

36. Ertl, P., Schuffenhauer, A., Renner, S., & Wegscheid-Gerlach, C. (2009). Fast calculation of molecular polar surface area as a sum of fragment-based contributions and its application to the prediction of drug transport properties. Journal of Medicinal Chemistry, 52(4), 1146–1154. 10.1021/jm8014425

37. Ciardiello, F., & Tortora, G. (2008). EGFR antagonists in cancer treatment. The New England Journal of Medicine, 358(11), 1160–1174. 10.1056/NEJMra0707704

38. Hochreiter, S., & Schmidhuber, J. (1997). Long short-term memory. Neural Computation, 9(8), 1735–1780. 10.1162/neco.1997.9.8.1735

39. Greff, K., Srivastava, R. K., Koutník, J., Steunebrink, B. R., & Schmidhuber, J. (2017). LSTM: A search space odyssey. IEEE Transactions on Neural Networks and Learning Systems, 28(10), 2222–2232. 10.1109/TNNLS.2016.2582924

40. Gers, F. A., Schraudolph, N. N., & Schmidhuber, J. (2002). Learning precise timing with LSTM recurrent networks. Journal of Machine Learning Research, 3, 115–143. https://www.jmlr.org/papers/volume3/gers02a/gers02a.pdf

41. Graves, A., Mohamed, A., & Hinton, G. (2013). Speech recognition with deep recurrent neural networks. 2013 IEEE International Conference on Acoustics, Speech and Signal Processing, 6645–6649. 10.1109/ICASSP.2013.6638947

42. Chung, J., Gulcehre, C., Cho, K., & Bengio, Y. (2014). Empirical evaluation of gated recurrent neural networks on sequence modeling. arXiv preprint arXiv:1412.3555. https://arxiv.org/abs/1412.3555

43. Sutskever, I., Vinyals, O., & Le, Q. V. (2014). Sequence to sequence learning with neural networks. Advances in Neural Information Processing Systems, 27, 3104–3112. https://papers.nips.cc/paper/2014/file/a14ac55a4f27472c5d894ec1c3c743d2-Paper.pdf

44. Segler, M. H., Kogej, T., Tyrchan, C., & Waller, M. P. (2018). Generating focused molecule libraries for drug discovery with recurrent neural networks. ACS Central Science, 4(1), 120–131. 10.1021/acscentsci.7b00512

45. Bjerrum, E. J. (2017). SMILES enumeration as data augmentation for neural network modeling of molecules. arXiv preprint arXiv:1703.07076. https://arxiv.org/abs/1703.07076

46. Polykovskiy, D., Zhebrak, A., Sanchez-Lengeling, B., Golovanov, S., Tatanov, O., Belyaev, S., & Kadurin, A. (2018). Molecular sets (MOSES): A benchmarking platform for molecular generation models. Frontiers in Pharmacology, 9, 5. 10.3389/fphar.2018.01135

47. Sutskever, I., Vinyals, O., & Le, Q. V. (2014). Sequence to sequence learning with neural networks. Advances in Neural Information Processing Systems, 27, 3104–3112. https://papers.nips.cc/paper/2014/file/a14ac55a4f27472c5d894ec1c3c743d2-Paper.pdf

48. Ciardiello, F., & Tortora, G. (2008). EGFR antagonists in cancer treatment. The New England Journal of Medicine, 358(11), 1160–1174. 10.1056/NEJMra0707704

49. Seshacharyulu, P., Ponnusamy, M. P., Haridas, D., Jain, M., Ganti, A. K., & Batra, S. K. (2012). Targeting the EGFR signaling pathway in cancer therapy. Expert Opinion on Therapeutic Targets, 16(1), 15–31. 10.1517/14728222.2011.645805

50. Normanno, N., De Luca, A., Bianco, C., Strizzi, L., Mancino, M., Maiello, M. R., & Salomon, D. S. (2006). Epidermal growth factor receptor (EGFR) signaling in cancer. Gene, 366(1), 2–16. 10.1016/j.gene.2005.10.018

51. Stamos, J., Sliwkowski, M. X., & Eigenbrot, C. (2002). Structure of the Epidermal Growth Factor Receptor kinase domain alone and in complex with a 4-anilinoquinazoline inhibitor. Journal of Biological Chemistry, 277(48), 46265–46272. 10.1074/jbc.M207135200

52. Bickerton, G. R., Paolini, G. V., Besnard, J., Muresan, S., & Hopkins, A. L. (2012). Quantifying the chemical beauty of drugs. Nature Chemistry, 4(2), 90–98. 10.1038/nchem.1243

